# Dishevelled 2 coordinates aortic Notch signaling through primitive myeloid and hemogenic endothelial cells during hematopoietic stem cell specification

**DOI:** 10.64898/2026.09.29.755325

**Authors:** Kelsey A. Carpenter, Amber D. Ide, Jacquelyn Jacobs, Emma Rudisel, Nina Sheely, Margarita M. Parada-Kusz, Stephanie Grainger

## Abstract

Hematopoietic stem cells (HSCs) arise from the hemogenic endothelium in the dorsal aorta through the endothelial-to-hematopoietic transition, a tightly regulated developmental process requiring coordinated signaling from the developing hematopoietic niche. Wnt and Notch signaling are essential for HSC specification, with Wnt functioning upstream of Notch; however, the molecular mechanisms by which these pathways are coordinated during HSC development remain poorly understood. Here, we identify Disheveled 2 (Dvl2), a key intracellular Wnt scaffold protein, as an essential regulator of HSC identity specification in zebrafish. Loss of *dvl2* impaired HSC specification and reduced Notch signaling in the dorsal aorta, despite normal canonical Wnt activity in the developing endothelium, suggesting that Dvl2 regulates HSC specification independently of endothelial β-catenin signaling. Hemogenic endothelial-specific expression of the Notch intracellular domain rescued HSC defects in *dvl2* morphants, demonstrating that Dvl2 functions upstream of Notch activation in the hemogenic endothelium during HSC specification. In addition, loss of *dvl2* reduced primitive macrophages and neutrophil populations. Restoration of primitive myeloid cells through *gata1* knockdown rescued both endothelial Notch signaling and HSC defects, demonstrating that Dvl2 is also required for the supportive immune cells that induce HSC identity. Together, these findings reveal that Dvl2 coordinates hemogenic endothelial and immune niche cues to control HSC identity specification.

## INTRODUCTION

Hematopoiesis occurs through two sequential and highly conserved developmental waves to establish the vertebrate blood system. Primitive hematopoiesis generates transient embryonic erythrocytes, macrophages and neutrophils, cells required for early oxygen transport and innate immunity for embryonic survival^1,2^. During the second, definitive, wave of hematopoiesis, hematopoietic stem cells (HSCs) capable of self-renewal and multilineage differentiation to sustain lifelong blood production are generated^3,4^. During definitive hematopoiesis, HSCs arise from specialized hemogenic endothelial cells located in the ventral wall of the dorsal aorta through the endothelial-to-hematopoietic transition (EHT), during which endothelial cells acquire hematopoietic identity and bud into circulation as nascent HSCs^5–7^. Because HSCs emerge in limited numbers, their specification and emergence require precise coordination of developmental signaling pathways, disruption of which can impair hematopoiesis and contribute to hematopoietic malignancies^8–13^.

Wnt signaling is required for HSC specification, proliferation, maintenance and differentiation^12,14–25^. In general, a secreted Wnt ligand engages Frizzled receptors and co-receptors, promoting recruitment of the intracellular scaffold Disheveled (Dvl) near the plasma membrane, where it participates in downstream canonical (β-catenin dependent) and non-canonical (β-catenin independent) Wnt signaling^26–33^. Beyond their established role in Wnt signaling, Dvl proteins also function as multifunctional signaling hubs that regulate protein trafficking, cell polarity, cytoskeletal organization and signal pathway crosstalk in a context-dependent manner^34–38^. Mammals express three *DVL* paralogs (*DVL1-3*), with both overlapping and distinct functions and altered *DVL* expression has been associated with blood cancers^38–40^. Whether individual Dvl proteins coordinate distinct developmental signaling networks during HSC development remains unknown.

Notch signaling is also essential for HSC specification through a series of signals promote arterial identity, hemogenic endothelial specification and EHT^41–45^. Notch receptors are engaged by ligands of the Delta/Serrated/LAG-2 family, which leads to intracellular proteolytic Notch receptor cleavage to generate a Notch Intracellular Domain (NICD), which is released and translocated to the nucleus where it acts as a transcriptional activator^46–48^. Several signaling pathways operate upstream of Notch activation during HSC specification^8,49–52^. For example, non-canonical Wnt16 signaling regulates expression of Notch ligands *deltaC* and *deltaD* in the somites, establishing a signaling environment required for aortic Notch activation and HSC specification^8,44^. In addition to these direct developmental signals, HSC specification is shaped by the primitive immune microenvironment. For example, once primitive macrophages and neutrophils migrate to the dorsal aorta, they release inflammatory factors required to activate Notch signaling to promote EHT ^53–55^. Thus, Notch signaling during HSC specification is subject to convergent regulation through multiple niche cues, including the somites and immune cells.

Here, we identify Dvl2 as a regulator of hematopoiesis during zebrafish development. Loss of *dvl2* impairs HSC specification through two mechanisms that converge on Notch signal activation in the aorta. In the hemogenic endothelium, loss of *dvl2* reduces expression of the Notch components *jag1a* and *adam10a*, while overactivation of Notch signaling in these cells rescues HSC development, demonstrating that Dvl2 functions upstream of Notch. In parallel, loss of *dvl2* reduces primitive myeloid populations and restoring these populations rescues Notch signaling and HSC specification, demonstrating that the myeloid defect contributes to impaired Notch signaling through an extrinsic mechanism. Our findings identify Dvl2 as a regulator that impacts both the hemogenic endothelium and its surrounding niche to sustain Notch activity during HSC specification.

## RESULTS

### Dvl2 is required for HSC specification

There are numerous signaling cues upstream of aortic Notch activation, including Wnt signaling. The *dvl* gene family encodes scaffolding proteins required for Wnt signaling and other processes^37,38^. The zebrafish genome encodes five Dvl paralogs (*dvl1a, dvl1b, dvl2, dvl3a, dvl3b*), which exhibit both overlapping and distinct functions during embryonic development^56,57^; however, which of these are required for Notch activation and HSC specification are unknown. HSC emergence through the endothelial-to-hematopoietic transition begins around 26 hours post fertilization (hpf), marked by the expression of HSC-associated genes such as *gata2b* and *runx1*^6–8,45,58^. However, the inductive cues that drive HSC identity specification occur earlier than 26 hpf, as endothelial cells migrate from the region lateral to the somites to the midline^6,8,9,45,58,59^. To identify candidate *dvls* required for HSC specification, we analyzed publicly available single-cell RNA-sequencing data from DanioCell, focusing on endothelial populations at 20 hpf^60,61^. This analysis revealed that *dvl2* and *dvl3a* are the predominant Dvl paralogs expressed in endothelial cells at 20 hpf (Fig. S1A), identifying them as candidate regulators of HSC specification.

To investigate the requirement for *dvl2* and *dvl3a* in HSC specification, we performed loss-of-function experiments using both ATG- and splice-blocking morpholinos (MOs) targeting *dvl2* or *dvl3a* and assessed the formation of *runx1+* HSCs at 26 hpf using whole-mount *in situ* hybridization (WISH). Knockdown of *dvl2* (Fig. 1A, S1B), but not *dvl3a* (Fig. S1C), led to a reduction in *runx1+* cells in the dorsal aorta at 26 hpf, suggesting an impact on HSC specification. We observed a similar loss of *runx1:eGFP;kdrl:mCherryNLS* double positive HSCs in the dorsal aorta in *dvl2* morphant animals (Fig. 2B). To validate the specificity of our *dvl2* MO phenotype, we generated a CRISPR/Cas9-mediated mutant targeting the proximal promoter and translational start site of *dvl2*, producing a 473 bp deletion predicted to yield a null allele (hereafter referred to as *dvl2^-/-^*) (Fig. S1D). Consistent with the MO results, *dvl2^-/-^* embryos had fewer *runx1+* cells in the dorsal aorta (Fig. 1C), supporting the specificity of the MO phenotype.

**Figure 1.**
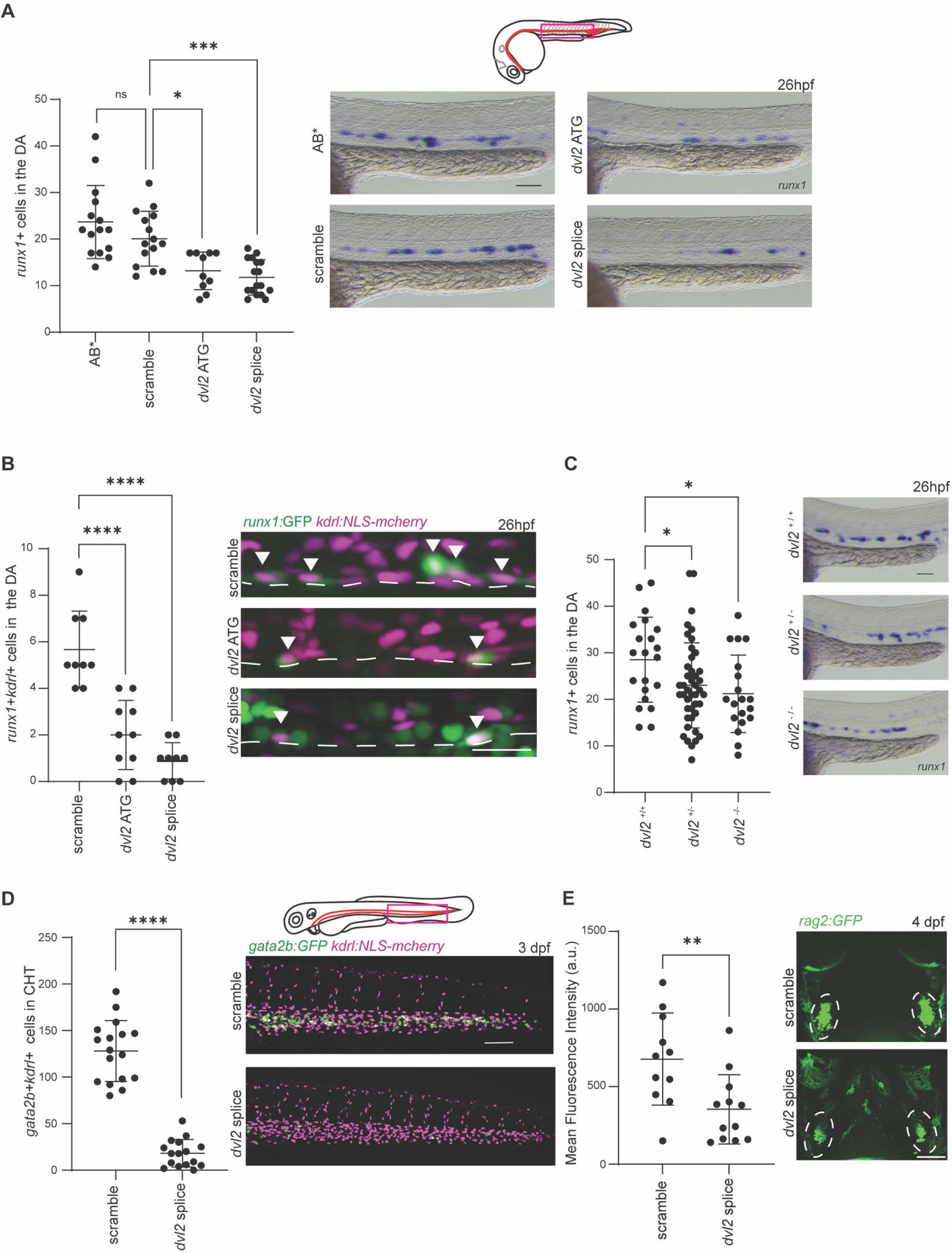
Dvl2 is required for HSC specification. **A.** Wild-type (AB*) embryos were injected with scramble control MO, *dvl2* ATG MO or splice blocking MO, embryos fixed at 26 hpf and analyzed by WISH for *runx1*. Representative images of *runx1+* cells within the dorsal aorta (boxed region) and quantification of *runx1+* cells are shown. Each dot represents one biological replicate. One-Way ANOVA with post hoc Tukey comparisons; * p<0.05, *** p<0.001. Scale bar = 100 µm. **B.** *runx1:GFP;kdrl:NLS-mcherry* embryos were injected with scramble MO, *dvl2* ATG MO or *dvl2* splice MO and imaged at 26 hpf. Representative confocal images and quantification of *runx1+mcherry+* double positive HSCs (white arrowheads) in the bottom of the dorsal aorta (dashed line). Each dot represents a biological replicate, One-Way ANOVA with post hoc Tukey comparisons; **** p<0.0001. Scale bar = 25 µm. **C.** CRISPR/Cas9-generated *dvl2* mutant embryos were analyzed for *runx1+* cells by WISH at 26 hpf. Representative images and quantification for *runx1*+ cells within the DA. Each dot represents one biological replicate. One-Way ANOVA with post hoc Tukey comparisons; * p<0.05. Scale bar = 100 µm. **D.** *gata2b:GFP;kdrl:NLS-mcherry* embryos were injected with scramble MO or *dvl2* splice MO and imaged at 3 dpf. Representative confocal images and quantification of *gata2b+mcherry+* double positive HSPCs in the CHT (boxed region). Scale bar = 25 µm. Each dot represents a biological replicate, Two-tailed Student’s, **** p<0.0001. **E.** *Rag2:GFP* embryos were injected with scramble MO or *dvl2* MO and imaged at 4 dpf. Quantification of the mean *GFP* fluorescence intensity within the thymus, with each dot represents a biological replicate; Two-Tailed Student’s, ** p<0.01. Representative images of the thymus (outlined by white dotted circles), scale bar = 100 µm.

**Figure 2.**
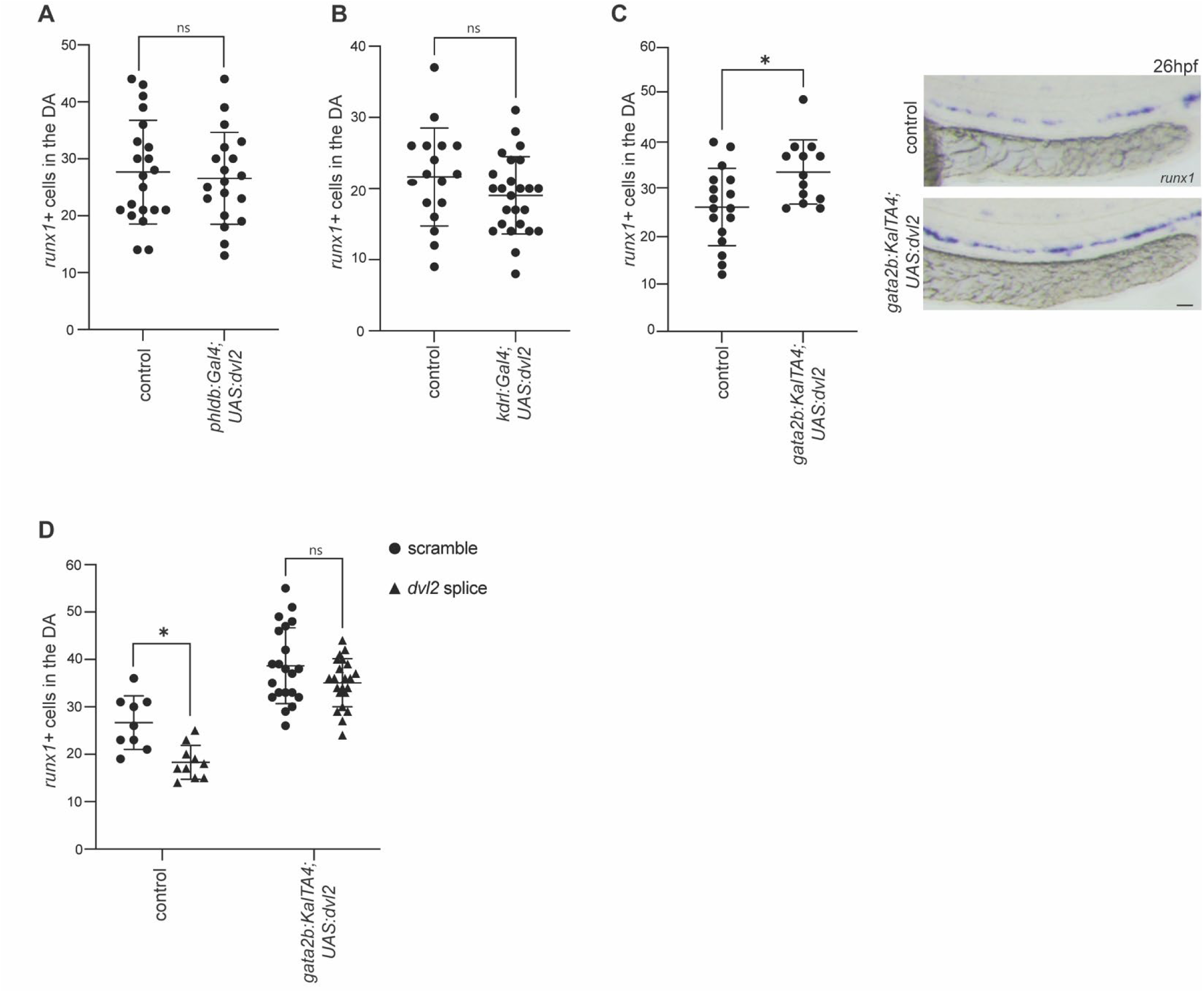
Dvl2 functions within the hemogenic endothelium to promote HSC specification. **A.** *phldb:Gal4* fish were crossed to *UAS:dvl2* fish, embryos fixed at 26 hpf and analyzed by WISH for *runx1*. Non-transgenic siblings served as controls. Quantification of *runx1+* HSPCs in the dorsal aorta is shown where each dot represents a biological replicate; Two-tailed Student’s. **B**. *kdrl:Gal4* fish were crossed to *UAS:dvl2* fish, embryos were fixed at 26 hpf and analyzed by WISH for *runx1*. Non-transgenic siblings served as controls. Quantification of *runx1+* HSPCs in the dorsal aorta where each dot represents a biological replicate; Two-tailed Student’s. **C.** *gata2b:KalTA4* fish were crossed to *UAS:dvl2* fish, embryos were fixed at 26 hpf and analyzed by WISH for *runx1*. Non-transgenic siblings served as controls. Quantification of *runx1+* HSPCs in the dorsal aorta where each dot represents a biological replicate. Two-tailed Student’s; * p< 0.05. Representative WISH images of control and *gata2b:KalTA4;UAS:dvl2* embryos are shown, scale bar = 50 µm **C.** *gata2b:KalTA4;UAS:dvl2* embryos were injected with scramble MO or *dvl2* splice MO, fixed at 26 hpf and analyzed by WISH for *runx1*. Non-transgenic sibling embryos served as controls. Quantification of *runx1+* HSPCs in the dorsal aorta where each dot represents a biological replicate. One-Way ANOVA with post hoc Tukey comparisons; * p <0.05.

Following specification and emergence from the aorta, HSCs migrate to and expand in the caudal hematopoietic tissue (CHT), a secondary niche similar to the fetal liver in mammals^5,62^. To determine whether the early HSC specification defect observed upon *dvl2* knockdown persists at later stages, we analyzed *gata2b:GFP;kdrl:mCherryNLS* embryos at 3 days post fertilization (dpf). Consistent with our earlier findings, *dvl2* morphants exhibited a reduction in *gata2b+kdrl+* HSCs in the CHT (Fig. 1D), indicating that the defect in HSC development is sustained beyond initial specification. We next asked whether this defect extends to downstream lymphoid development by examining thymocyte formation, a later stage of hematopoietic differentiation, using *rag2:GFP* reporter embryos. Thymocyte expression was reduced in the thymus in the *dvl2* morphants (Fig. 1E), consistent with impaired development of early lymphoid progenitors. Furthermore, morphant animals exhibited normal mesoderm development (*tbx20*, Fig. S2A and *hand2* Fig. S2B), somites (*myod*, Fig. S2C), primitive blood (*gata1*, Fig. S2D), vasculature (*kdrl*, Fig. S2E) and arterial (*efnb2a*, Fig. S2F) suggesting that the HSPC defects are specific. Together, these data identify *dvl2* as a regulator of HSC specification in zebrafish.

### Dvl2 plays a tissue-specific role during HSC development

To investigate the tissue-specific requirements for Dvl2 during HSC development, we generated *UAS:dvl2* transgenic animals that enable *Gal4*-dependent expression of *dvl2* in distinct cell populations. Somitic Wnt16 is required for HSC specification; therefore, we overexpressed *dvl2* in the developing somites using *phldb1:Gal4;UAS:dvl2* animals and found that *runx1+* HSCs in the dorsal aorta at 26 hpf were unchanged (Fig. 2A), indicating that somite-derived *dvl2* does not affect HSC specification^8^. Because *dvl2* is expressed in endothelial cells during the Wnt-responsive window of HSC development prior to 20 hpf (Fig. S1A), we examined whether endothelial overexpression of *dvl2* could influence HSC specification. In *kdrl:Gal4;UAS:dvl2* embryos, endothelial-specific expression of *dvl2* did not alter HSC numbers (Fig. 2B), suggesting that broad endothelial overexpression is insufficient to affect specification. We next tested whether *dvl2* overexpression within the hemogenic endothelium could modulate HSC development. In *gata2b:KalTA4;UAS:dvl2* animals, we observed a significant increase in HSCs in the dorsal aorta (Fig. 2C), indicating that *dvl2* can function in hemogenic endothelial cells to promote HSC specification. To further validate that Dvl2 functions within the hemogenic endothelium, we injected *dvl2* MO into *gata2b:KalTA4;UAS:dvl2* embryos to determine whether hemogenic endothelial expression of *dvl2* was sufficient to rescue the MO-induced phenotype. Knockdown of *dvl2* in *gata2b:KalTA4;UAS:dvl2* embryos increased HSPCs in the dorsal aorta (Fig 2D). Together, these data suggest that Dvl2 in the hemogenic endothelium is required for HSC specification.

### Dvl2 modulates Notch signaling in the developing endothelium

Multiple Notch signaling inputs across distinct tissues are essential for HSC specification, and promoting Notch activity in the floor of the dorsal aorta is the penultimate event required to inititate *runx1* expression^9,44,45,58^. Non-canonical Wnt16 is required upstream of this activation, through promoting Notch ligands expression^8,9^. This is in contrast to canonical Wnt signaling, which has been implicated in the subsequent expansion of HSCs, a process that occurs downstream of Notch activation^17,18^. To determine if Dvl2 acts upstream of aortic Notch activation, we analyzed *tp1:GFP;kdrl:NLS-mCherry* animals, which have GFP expression driven by the Notch-responsive *tp1* element and nuclear mCherry driven in endothelial cells*. tp1:GFP* fluorescence reflects active Notch signaling in endothelial cells. We quantified GFP fluorescence in the endothelial cells lining the DA, where Notch activity is required for HSC specification, and found that *dvl2* knockdown reduced *tp1:GFP* fluorescence (Fig. 3A). Canonical Wnt signaling can be measured using *7XTCF:GFP* reporter fish, where GFP is driven by 7 tandem LEF/TCF response elements^63^. Injection of *dvl2* MO into *7XTCF:GFP;kdrl:mCherryNLS* embryos led to comparable GFP fluorescence in the endothelial populations (mCherry) compared to control morphants at 16.5 hpf (Fig. S3A-C), a timepoint when canonical Wnt signaling is known to be required for HSC amplification^17^, suggesting that canonical Wnt is not impacted in this context. We further confirmed this using qPCR for *axin2,* a canonical Wnt target gene^64^, which we found was unchanged by loss of *dvl2* at 26 hpf (Fig. S3D). Together, these results suggest that Dvl2 operates upstream of Notch signaling during HSC specification.

**Figure 3.**
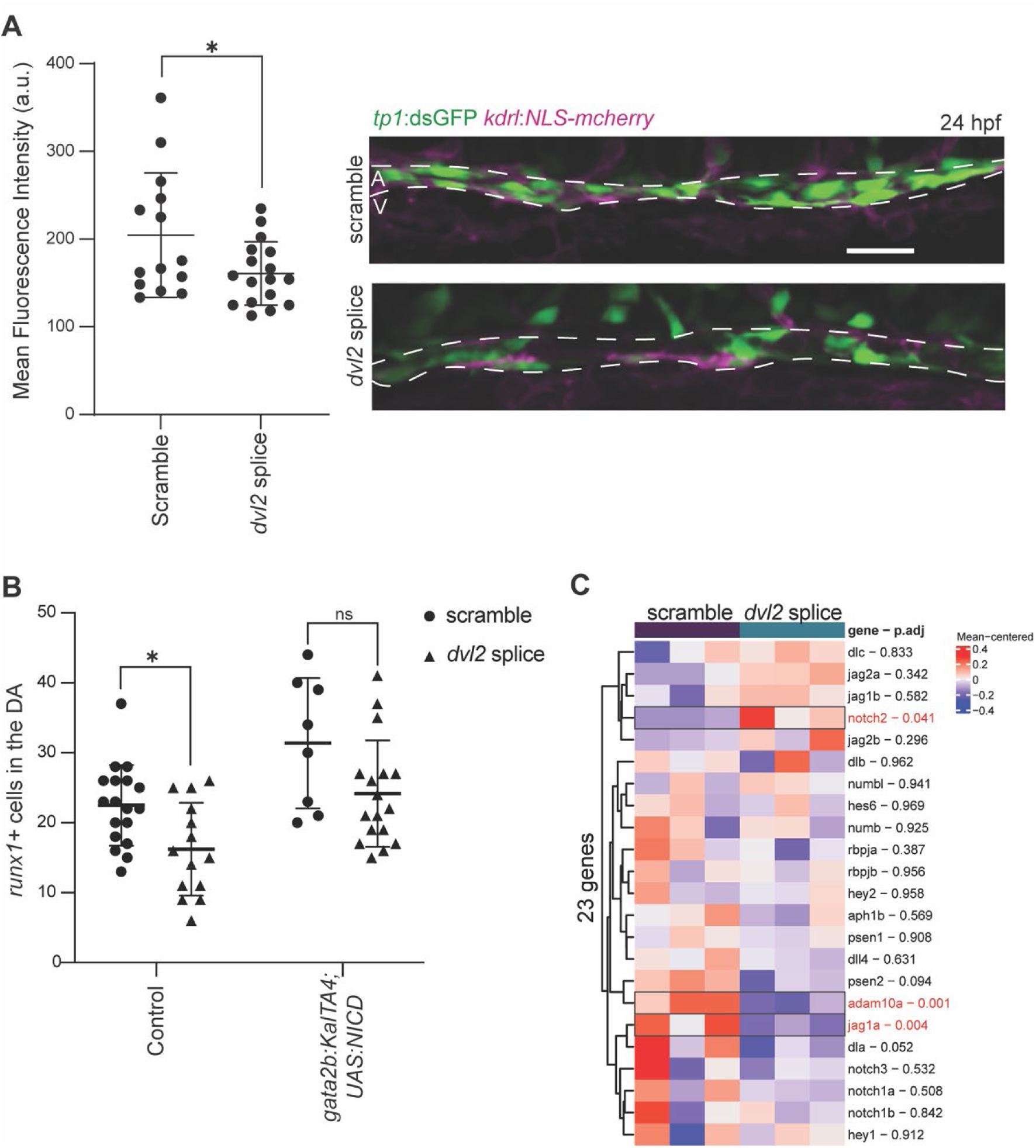
Dvl2 regulates Notch signaling in the hemogenic endothelium during HSC specification. **A.** *tp1:GFP* fish were crossed to *kdrl:NLS-mcherry* fish and injected with scramble MO or *dvl2* splice MO. Notch reporter activity was assessed by quantifying GFP fluorescence intensity within the dorsal aorta at 24 hpf. Each dot represents the average GFP fluorescence intensity from an individual embryo. Two-tailed Student’s, * p< 0.05. Representative images are shown, scale bar = 50 µm. **B.** *gata2b:KalTA4* fish were crossed to *UAS:NICD* fish and embryos were injected with scramble MO or *dvl2* splice MO, fixed at 26 hpf and analyzed by WISH for *runx1*. Non-transgenic embryos served as controls. Quantification of *runx1+* HSPCs in the dorsal aorta is shown, with each dot representing one biological replicate. One-Way ANOVA with post hoc Tukey comparisons; * p<0.05. **C.** Heatmap showing differential gene expression analysis of RNA-sequencing data from FACS-isolated *runx1+* cells from scramble MO and *dvl2* splice MO-injected embryos at 26 hpf. Genes with significantly altered expression are highlighted in red.

To determine whether impaired Notch signaling underlies the HSC defect observed in *dvl2* morphants, we restored Notch activity within the hemogenic endothelial cells using *gata2b:KalTA4*;*UAS:NICD* animals, which express the Notch Intracellular Domain (NICD) in hemogenic endothelial cells^65^. We observed that hemogenic endothelial expression of *NICD* was sufficient to restore *runx1+* HSCs observed in *dvl2* morphants (Fig. 3B), demonstrating that activation of Notch signaling in the hemogenic endothelium is downstream of Dvl2 in HSC specification.

To identify how Dvl2 regulates Notch signaling, we analyzed RNA-sequencing data from FACS-isolated *runx1+* cells collected from scramble- and *dvl2* MO-injected embryos at 26 hpf. Differential gene expression analysis revealed reduced expression of *jag1a*, a Notch ligand that promotes receptor activation^55^ and *adam10a*, which encodes the metalloprotease responsible for ligand-induced cleavage of the Notch receptor, an essential step preceding NICD release and downstream transcriptional activation^66^ (Fig. 3C). Reduced expression of both *jag1a* and *adam10a* would therefore be predicted to diminish Notch pathway activation. Consistent with this model, forced expression of NICD bypassed these upstream requirements and restored HSC specification in *dvl2*-defiecient embryos (Fig. 3B). Together, these findings indicate that Dvl2 functions upstream of Notch activation to promote HSC specification by maintaining expression of key components required for Notch receptor activation.

### Dvl2 regulates myeloid cell development

The Notch ligand *jag1a* is induced by inflammatory signaling from myeloid cells, with TNFa produced predominantly by neutrophils activating Tnfr2 in the aortic endothelium to promote *jag1a* expression and Notch signaling during HSPC emergence^55,67^. To assess the effect of *dvl2* loss on myeloid populations, we analyzed *mpeg:GFP* embryos and observed a reduction in GFP+ macrophages at 4 dpf in *dvl2* morphants (Fig. 4A). Similarly, there were also fewer mpx+ neutrophils in *dvl2* morphants (Fig. 3B), supporting a role for Dvl2 in primitive myelopoiesis. In contrast, erythropoiesis was unaffected, as *gata1:dsRed+* erythrocytes were unchanged in *dvl2* morphants (Fig. 4C), indicating that Dvl2 impacts myeloid lineages independent of erythroid cells.

**Figure 4.**
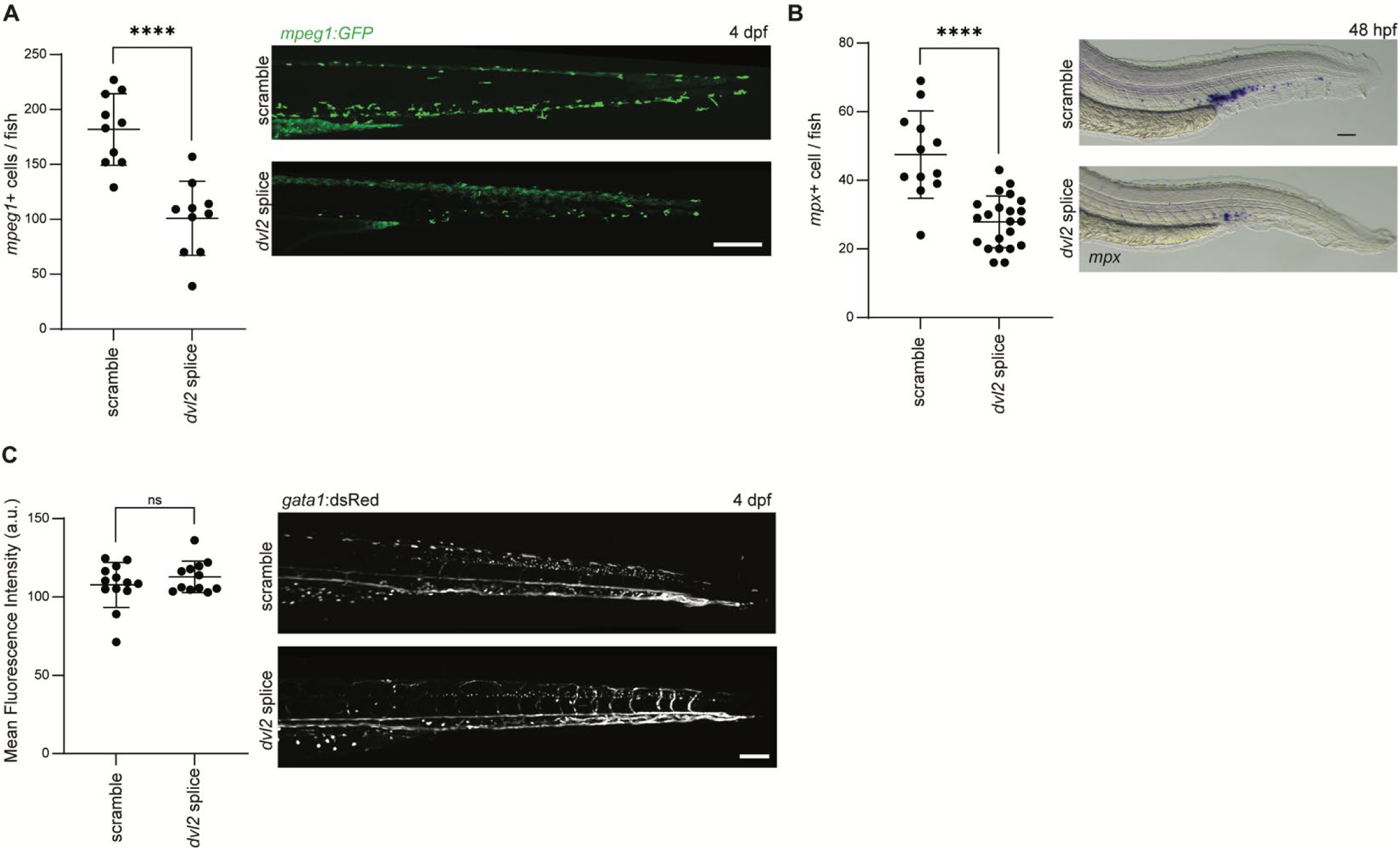
**Dvl2 regulates primitive hematopoiesis**. **A**. *mpeg1:GFP* embryos were injected with scramble MO or *dvl2* splice MO and imaged at 4 dpf. Quantification of *mpeg1:GFP+* macrophages in the CHT, with each dot represents one biological replicate; Two-tailed Student’s, **** p< 0.0001. Representative confocal Z-stacks images of the CHT, scale bar = 200 µm. **B.** AB* embryos were injected with scramble or *dvl2* splice MO, fixed at 48 hpf and analyzed by WISH for *mpx*. Quantification of *mpx+* neutrophils in the CHT is shown, with each dot representing one biological replicate; Two-tailed Student’s, **** p<0.0001. Representative images are shown, scale bar = 50 µm. **C.** *gata1:dsRed* embryos injected with scramble or *dvl2* splice MO and imaged at 4 dpf. Quantification of the mean *dsRed* fluorescent intensity in the CHT, with each dot represents one biological replicate; Two-Tailed Student’s. Representative images in the CHT are shown, scale bar = 100 µm.

### Myeloid cells restore endothelial Notch signaling and rescue HSC specification in *dvl2* morphants

Myeloid cells are an essential component of the developing hematopoietic niche: neutrophils promote HSC emergence through inflammatory signaling that enhances Notch activity in the DA^53,55^, whereas macrophages facilitate HSPC emergence by remodeling the extracellular matrix and supporting HSC emergence, clearance and proliferation^68,69^. Our findings led us to hypothesize that loss of macrophages and neutrophils in *dvl2* morphants contribute to the HSC defect. To test this hypothesis, we increased primitive myeloid output by knocking down *gata1a*, which shifts hematopoietic differentiation away from the erythroid lineage and toward myeloid fates^70^. We found that *dvl2* knockdown significantly reduced *mpeg:mCherry+* macrophages compared to scramble controls and co-injection of *gata1a* MO and *dvl2* splice MO increased macrophage numbers throughout the embryo (Fig. 5Ai, 5B), as well as near the dorsal aorta (Fig. 5Aii, 5C). Similarly, we found that *dvl2* knockdown significantly reduced *mpx:GFP+* neutrophils and increasing primitive myeloid output by co-injecting *gata1a* MO and *dvl2* splice MO increased neutrophils numbers in both the whole embryo (Fig. 5Di, 5E) and specifically near the dorsal aorta (Fig. 5Dii, 5F). Therefore, *gata1a* reduction was sufficient to increase the number of macrophages and neutrophils in *dvl2* morphants. Finally, we found that co-injection of *gata1a* MO and *dvl2* splice MO also restored *runx1+* HSCs in the dorsal aorta (Fig 5G), supporting that myeloid cells operate downstream of Dvl2 during HSC specification.

**Figure 5.**
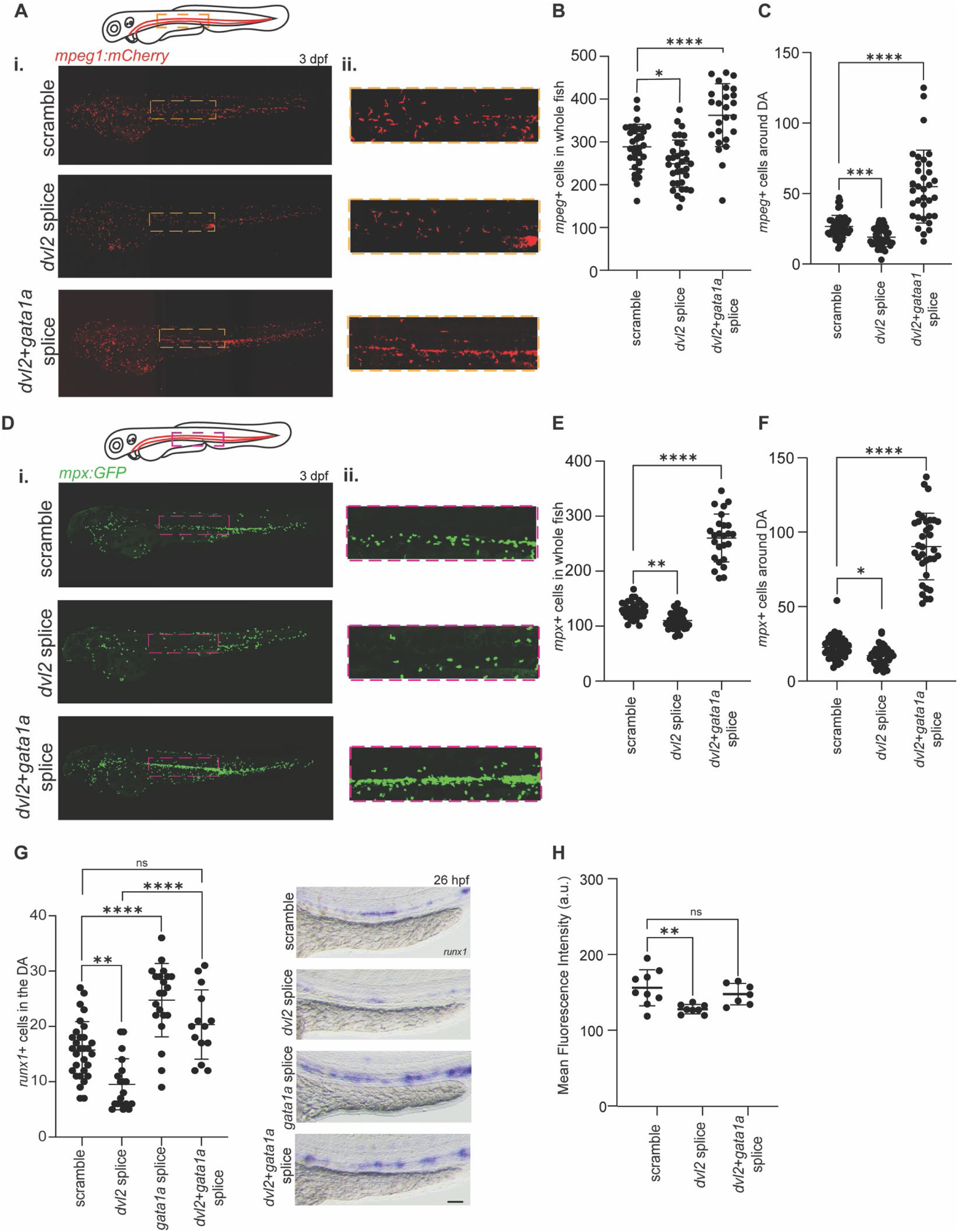
Primitive immune cells rescue HSC specification and Notch signaling in *dvl2* knockdown embryos. A-C. *mpeg1:mCherry;mpx:GFP* embryos were injected with scramble, *dvl2* or *dvl2+gata1a* MOs and imaged at 26 hpf. Representative images and quantification of the number of *mpeg:mCherry+* macrophages throughout the whole embryo (**Bi, C**) and near the dorsal aorta (**Bii, D**) are shown. Each dot represents a biological replicate; One-Way ANOVA with post hoc Tukey comparisons, * p<0.05, *** p< 0.001, **** p<0.0001. **D-F**. *mpeg1:mCherry;mpx:GFP* embryos were injected with scramble, *dvl2* or *dvl2+gata1a* MO and imaged at 26 hpf. Representative images and quantification of the number of *mpx:GFP+* neutrophils throughout the whole fish (**Ei, F**) and near the dorsal aorta (**Eii, G**) are shown. Each dot represents a biological replicate; One-Way ANOVA with post hoc Tukey comparisons, * p<0.05, ** p<0.01, **** p<0.0001. **G.** AB* embryos were injected with scramble, *dvl2*, *gata1a*, or *dvl2+gata1a* MOs, fixed at 26 hpf and analyzed by WISH for *runx1*. Quantification of *runx1+* HSCs in the dorsal aorta is shown, with each dot representing one biological replicate; One-Way ANOVA with post hoc Tukey comparisons, ** p<0.01, **** p<0.0001. Representative images are shown, scale bar = 125 µm. **H**. *tp1:dsGFP;kdrl:mCherry* embryos were injected with scramble, *dvl2* or *dvl2+gata1a* MOs and imaged at 24 hpf. Endothelial Notch reporter activity was assessed by quantifying GFP fluorescence intensity within the DA. Each dot represents the average GFP fluorescence intensity from an individual embryo. Two-tailed Student’s, * p< 0.05.

Primitive neutrophils promote HSC emergence by enhancing endothelial Notch signaling^55^. We assessed Notch activity using the *tpi1:dsGFP;kdrl:mCherry-NLS* in the context of *gata1a* and *dvl2* knockdown. Using this strategy, we found that increasing primitive myeloid populations through *gata1a* MO restored endothelial Notch reporter activity in *dvl2* morphant embryos within the dorsal aorta at 26 hpf (Fig. 5H). Thus, expansion of primitive myeloid cells rescues both endothelial Notch signaling and HSC specification following loss of *dvl2*.

## DISCUSSION

HSC specification requires the integration of signals in hemogenic endothelium with cues from the surrounding niche environments. Although Wnt and Notch signaling are both established regulators of HSC development, how these pathways are coordinated within and between tissues during the narrow developmental remains unclear. Here, we identify Dvl2 as a regulator of HSC specification that coordinates both endothelial-intrinsic and microenvironmental signals converging on Notch activity. Loss of *dvl2* reduced Notch activity and impaired HSC specification without detectable changes in canonical Wnt signaling. In the hemogenic endothelium, Dvl2 promotes the expression of Notch activators *jag1a* and *adam10a,* and constitutive Notch activation rescues the *dvl2* morphant HSC phenotype. In parallel, *dvl2* knockdown reduced macrophage and neutrophil populations, and restoring these cells rescued both endothelial Notch signaling and HSC specification. Together, these findings support a model in which Dvl2 couples hemogenic endothelial signaling with cues from the embryonic immune microenvironment to sustain Notch activation during HSC specification.

Our findings establish a cell-specific requirement for Dvl2 during HSC development. Although Dvl2 is broadly expressed in the developing endothelium, increasing Dvl2 specifically in hemogenic endothelial cells, not in the broader endothelium or somites, increased HSC numbers, indicating that Dvl2 activity is particularly important within cells that have received HSC specification cues, rather than reflecting a general vasculature or somitic requirement. This requirement appears to act upstream of Notch activation: *dvl2* loss reduced endothelial Notch reporter activity, while hemogenic endothelial expression of NICD restored HSC specification. This mechanism is distinct from the role for Dvl2 in canonical Wnt signaling, as *dvl2* loss did not alter endothelial 7XTCF activity or *axin2* expression during this window.

Transcriptional analysis further supports altered Notch pathway regulation following *dvl2* loss. Alongside reduced *jag1a* and *adam10a* expression, we observed increased in *notch2,* unlikely to reflect compensatory signaling, since Notch2 is dispensable for HSC specification in zebrafish and mice^71^. In zebrafish, Notch1a/b, not Notch2, act in the hemogenic endothelium to drive HSC specification^44^.

Because the expression of these required Notch receptors was not reduced following *dvl2* loss, the decrease in Notch activity likely does not stem from reduced receptor expression. Instead, reduced expression of *jag1a* and *adam10a* diminished Notch reporter activity, and rescue by NICD together suggest that Dvl2 promotes Notch activation through regulation of other pathway components, acting at the level of pathway activation rather than receptor transcription.

These findings diverge from the classical view of Dvl2 as a Wnt mediator. Previous work in zebrafish showed that non-canonical Wnt16 signaling regulates somite-derived Notch ligands to drive HSC specification^8^, whereas canonical Wnt signaling acts later to promote HSC expansion and maintenance^14,17,18^. Our analyses focused on the developmental window preceding EHT, when canonical Wnt is thought to play a minor role; thus, our findings do not exclude additional Dvl functions in canonical Wnt signaling at later stages, but instead reveal an earlier, β-catenin-independent requirement. This is consistent with reports in *Drosophila* that Dvl genetically interacts with Notch and Delta and physically associates with the NICD, suggesting Dvl can modulate Notch signaling independently of canonical Wnt signaling^72^. Whether Dvl2 physically associates with Notch pathway components and whether this regulates receptor processing or ligand activity, will be important to determine.

In addition to its role in the hemogenic endothelium, our findings reveal an additional role for Dvl2 in establishing the myeloid environment that supports HSC specification. Macrophages and neutrophils are recognized as active components of the embryonic hematopoietic niche rather than simply transient blood populations. Neutrophils can promote HSC emergence through inflammatory signaling that enhances endothelial Notch activity^55^, while macrophages contribute to the developing niche through extracellular matrix remodeling and regulation of HSC emergence and proliferation^68^. We found that loss of *dvl2* reduced both macrophages and neutrophils, thus, the hematopoietic phenotype caused by loss of *dvl2* extends beyond the hemogenic endothelium and includes disruption of the cellular microenvironment surrounding emerging HSCs.

The functional relationship between these myeloid defects and impaired HSC specification was demonstrated by increasing output of both macrophages and neutrophils in *dvl2* morphants. These findings indicate that the reduction of myeloid cells is not simply a consequence of impaired hematopoiesis but contributes functionally to the HSC phenotype. Given that primitive neutrophils promote Notch activation through inflammatory signaling, one possibility is that Dvl2 maintains a myeloid-derived signaling that reinforces *jag1a* expression and Notch activity within the DA. Lineage-specific manipulation will be required to distinguish whether these populations provide distinct signals or function cooperatively to support HSC development.

Collectively, our study identifies Dvl2 as a regulator of HSC specification that coordinates intrinsic and extrinsic developmental signals that converge through Notch activity. These findings expand the function of Dvl2 beyond its canonical role in Wnt signaling and demonstrate how a multifunctional signaling scaffold can integrate signals within hemogenic endothelium with cues from the surrounding immune microenvironment. More broadly, our work highlights the importance of coordinating endothelial signaling with the development cellular niche during HSC specification and provides a framework for understanding how multiple developmental signals converge to regulate the emergence of definitive hematopoiesis.

## ACKNOWLEDGEMENTS

Imaging was performed in part in the Van Andel Institute Optical Imaging Core (RRID:SCR_021968). Flow cytometry was performed with assistance from the Van Andel Flow Cytometry Core (RRID:SCR_022685). Vivarium staff (RRID:SCR_023211) are thanked for animal husbandry. Research reported in this publication was supported by the National Institute of General Medical Science under Award Number R35GM142779 (SG), by the American Cancer Society under Award Number PF-23-1037956-01-CCB (KAC) and by the National Institute of Cancer under Award Number T32CA251066 (ADI). The content is solely the responsibility of the authors and does not necessarily represent the official views of the National Institutes of Health or American Cancer Society.

## AUTHOR CONTRIBUTIONS

KAC, ADI, JJ, ER, NS and MRK designed, conducted and analyzed experiments, and edited the manuscript. KAC, ADI and SG conceived, designed, supervised experiments and analysis, and wrote the manuscript.

## METHODS

### Animals

Zebrafish were housed and maintained according to Van Andel Institute and local Institutional Animal Care and Use Committee policies. AB* zebrafish were used as wild-type animals in all experiments. The *Tg(runx1:eGFP)^y509Tg 73^, Tg(gata2b:KalTA4)^sd32Tg^* ^58^, *Tg(UAS:GFP)^mu^*^271^ ^74^, *Tg(kdrl:NLS-mCherry)^y173Tg^* ^75^, *Tg*(*7XTCF-X.laveis-siamois:eGFP)^ia4^* ^76^, *Tg(TP1:dsGFP), Tg(mpeg1:mCherry)^g122Tg^* ^77^, Tg(mpx:GFP)^i114^ ^78^, *Tg(gata1:dsRed)^sd2Tg^* ^79^, *Tg(Rag2:GFP), Tg(kdrl:Gal4)^bw9Tg^* ^44^*, Tg(UAS:myc:notch1a-intra)^kca^*^3^ ^65^ and *Tg(phldb1:KalTA4)^hzm7Et^* ^9^ lines have been previously described. All MOs were acquired from GeneTools and have the following sequences: ATG MO for *dvl2*: 5’-GCCATGTCTCTCAACCCTGCTAAAC-3’, splice-blocking MO for *dvl2*: 5’-TCACCACCCTGAGACACACAATTATCA-3’, ATG MO for *dvl3a*: 5’-AACTTTAGTCTCCCCCATTGCAGAC-3’, splice-blocking MO for *gata1a*: 5’-GTTTGGACTCACCTGGACTGTGTCT-3”, and scramble MO: 5’-CCTCTTACCTCAGTTACAATTTATA-3’. 1-cell stage zygotes were injected with 1.0 ng/nL *dvl2* MO, 1.69 ng/nL *gata1* MO, 1.0 ng/nL *dvl2* MO co-injected with 1.69 ng/nL *gata1a* MO, or 1.0 ng/nL standard MO, unless otherwise indicated in figures. Embryos and larvae were cultured to the ages indicated in figures in Essential 3 (E3) medium (5 mM NaCl, 0.17 mM KCl, 0.33 mM CaCl2, 0.33 mM MgSO4, 10^-5^ % Methylene Blue).

Clustered regularly interspaced short palindromic repeats (CRISPR)–Cas9 was used to generate germline mutants for *dvl2.* We used a previously established method of selecting two single guide RNAs based on their ability to cleave DNA *in vitro*. *Cas9* mRNA (Trilink, 100 ng) and 100ng of each single guide RNA (5’-AGTCCATACGAGTGTTTCTG – 3’ and 5’ – GACAAGGACACTGGTCTGGG – 3’) were used to achieve a 473 bp deletion, including the proximal promotor and ATG start codon of *dvl2*. Polymerase chain reaction (PCR) was used to genotype zebrafish and validate the deletion was completed using specific primers (Supplementary Table 1). These *dvl2* knock-out fish are referred to as *dvl2^-/-^*. Mutations were confirmed by sequencing F1 generation zebrafish. Lines are available upon request.

Dvl2 (ENSDARG00000056184.8) was PCR amplified from zebrafish cDNA. The transgenic plasmid for *UAS:dvl2-BFP;cmlc2:GFP* (referred to in the text as *UAS:dvl2)* was generated by inserting *dvl2* cDNA downstream of 4X tandem UAS in construct with *cmlc2:GFP* and Tol2 recombination sites in the backbone, and sequence validated by full-plasmid sequencing. *Tg(UAS:dvl2)* founders were established by injecting 25 pg of the *UAS:dvl2* generated plasmid with 100 pg transposase mRNA from the Tol2 kit at the one-cell stage^80^. The resultant animals were screened for GFP+ hearts and outcrossed to AB to establish germline founders.

### Whole mount *in situ* hybridization (WISH)

Whole mount *in situ* hybridization (WISH) protocols and probes for *runx1*, *cmyb, tbx20, hand2, myod, gata1, flk, rag1, efnb2a* and *mpx* have been previously described^8,9,82^. For WISH quantification experiments, multiple independent researchers were involved in cell number quantification (*runx1, cmyb, mpx*) to ensure reproducibility and minimize bias.

### qPCR

Total RNA was extracted from embryos at 26 hpf using a Quick-RNA Lysis Kit (Zymo Research) and cDNA was reverse transcribed using iScript Reverse Transcription Supermix (Bio-Rad). qPCR was performed using PowerUp SYBR Green Master Mix (Thermo Fisher Scientific) according to the manufacturer’s instructions, samples were run on a QuantStudio 6 Pro real-time PCR machine (Applied Biosystems), and analyzed using the 2^− ΔΔCt^ method^83^. Primers used are available as a Supplementary Table 1.

### Confocal imaging and analysis

Live or fixed larval zebrafish were embedded in 0.7% UltraPure LMP Agarose (16250) in glass bottom dishes and covered with 1X E3 media. Imaging was performed on the Andor Dragonfly 620-SR spinning disc mounted of a Leica DMi8 microscope equipped with 488, 561, and 640 nm lasers using 10X objective or 25X immersion objective.

*tp1:dsGFP;kdrl:NLS-mCherry* zebrafish were imaged at 24 hpf. Z-stack images were acquired using a 1 μm step size from 1 field of view that encompassed the entire DA. A ROI was set in the dorsal aorta region of each fish using Imaris analysis software (10.2.0) and mean fluorescence intensity of GFP was acquired. The brightness and contrast of all images were uniformly adjusted using histograms of intensity distributions to optimize visualization of fluorescence in representative images.

*Mpeg1:eGFP, rag2:GFP,* and *gata1:dsRed* zebrafish were imaged at 4 dpf. To capture the entire animal, Z-stack images were acquired from 6 fields of view and auto-stitched post-acquisition. The stitched 3D images were cropped to include a ROI from the end of the yolk extension to the end of the CHT. The ROI size was identical in all samples. The surface model on Imaris software was used to detect the number of fluorescent cells.

The *runx1:eGFP;kdrl:memCherry* and *gata2b:GFP;kdrl:NLS-mcherry* zebrafish were imaged at 3 dpf, respectively. Z-stack images were acquired using a 1 μm step size from 1 field of view that encompassed the entire DA. A ROI from the end of the yolk extension to the end of the CHT was analyzed and was identical for all samples. The spot analysis tool was used to identify and count GFP+ and memCherry+ double positive cells along the entirety of the dorsal aorta floor in all Z planes per image. The brightness and contrast of all images were uniformly adjusted using histograms of intensity distributions to optimize visualization of fluorescence in representative images.

Transgenic *mpeg1:mCherry* and *mpx:GFP* zebrafish lines were crossed and at 3 dpf Z-stack images were acquired with a 10X air objective using 20-µm optical stacks and maximum-intensity projections were generated for analysis. Cell quantification was performed in FIJI (ImageJ) using the "Find Maxima" function to assess total cell counts throughout the entire larva. To analyze the dorsal aorta region and ensure consistency across samples, a standardized region of interest (ROI) encompassing the dorsal aorta was applied to all images. All fluorescent cells within the defined ROI were quantified.

### Flow Cytometry and analysis

*Tg*(*7XTCF:eGFP;kdrl:NLS-mCherry)* zebrafish embryos were used for flow cytometry at 16.5 hpf. Embryos were manually dissociated in FACS Buffer (PBS p.H. 7.4, 2% FBS, 1mM EDTA) for each experimental condition and supplemented with 2 µg/mL DAPI to exclude dead cells. Single-cell suspensions were filtered through a 35 µm mesh and analyzed on a Beckman Coulter CytoFLEX S flow cytometer. GFP fluorescence was detected using a 488 nm laser and 525/40 bandpass filter and mCherry fluorescence was detected using a 561 nm laser and 610/20 bandpass filter. Mean fluorescence intensity was used to quantify GFP expression levels.

### RNA-Sequencing

*runx1:GFP* fish were injected with scramble MO or *dvl2* splice MO. For each biological replicate, six embryos were pooled (n= 3 biological replicates per condition). At 26 hpf, embryos were manually dissociated in FACS buffer (PBS containing 2% FBS, 1mM EDTA, 2 µg/mL DAPI), filtered through a 80 µm filter and GFP+ cells were isolated using a BD FACSAria cell sorter. Total RNA was extracted from sorted cells using a Quick-RNA Lysis Kit (Zymo Research).

Libraries were prepared by the Van Andel Genomics Core from 1.8 ng of total RNA using the Watchmaker RNA Library Prep kit (Watchmaker Genomics). Ribosomal RNA material was reduced using the QIAseq FastSelect – rRNA Fish Kit (Qiagen). RNA was sheared to 300-400 bp. Prior to PCR amplification, cDNA fragments were ligated to IDT for Illumina TruSeq UD Indexed adaptors (Integrated DNA Technologies). Quality and quantity of the finished libraries were assessed using a combination of Agilent DNA High Sensitivity chip (Agilent Technologies, Inc.) and QuantiFluor dsDNA System (Promega Corp.). Individually indexed libraries were pooled and 75 bp, paired end sequencing was performed on an Element AVITI sequencer using a high-output, 150 bp sequencing kit (Element Biosciences) to an average depth of 71M reads per sample. Base calling was performed on instrument and AVITI OS (v2.6.2) output was demultiplexed and converted to FastQ format with Element Biosciences Bases2fastq (v2.3.0). Adaptor sequences and low-quality bases were trimmed from raw sequencing reads using Trim Galore^84^. Trimmed reads were aligned to the zebrafish reference genome (GRCz11.115) using STAR with the “-quantMode GeneCounts” option enabled to generate gene-level count matrices. Differential gene expression analysis was performed using DESeq2^85,86^. Raw gene counts were normalized, dispersion estimates were calculated and differential expression between scramble MO and *dvl2* MO-treated samples was assessed using a generalized linear model. P-values were adjusted for multiple hypothesis testing using the Benjamini-Hochberg method to control the false discovery rate (FDR). Gene set enrichment analysis and pathway visualization were performed using the R Bioconductor package, clusterProfiler^87–89^.

---

**Supplementary Figure 1.**
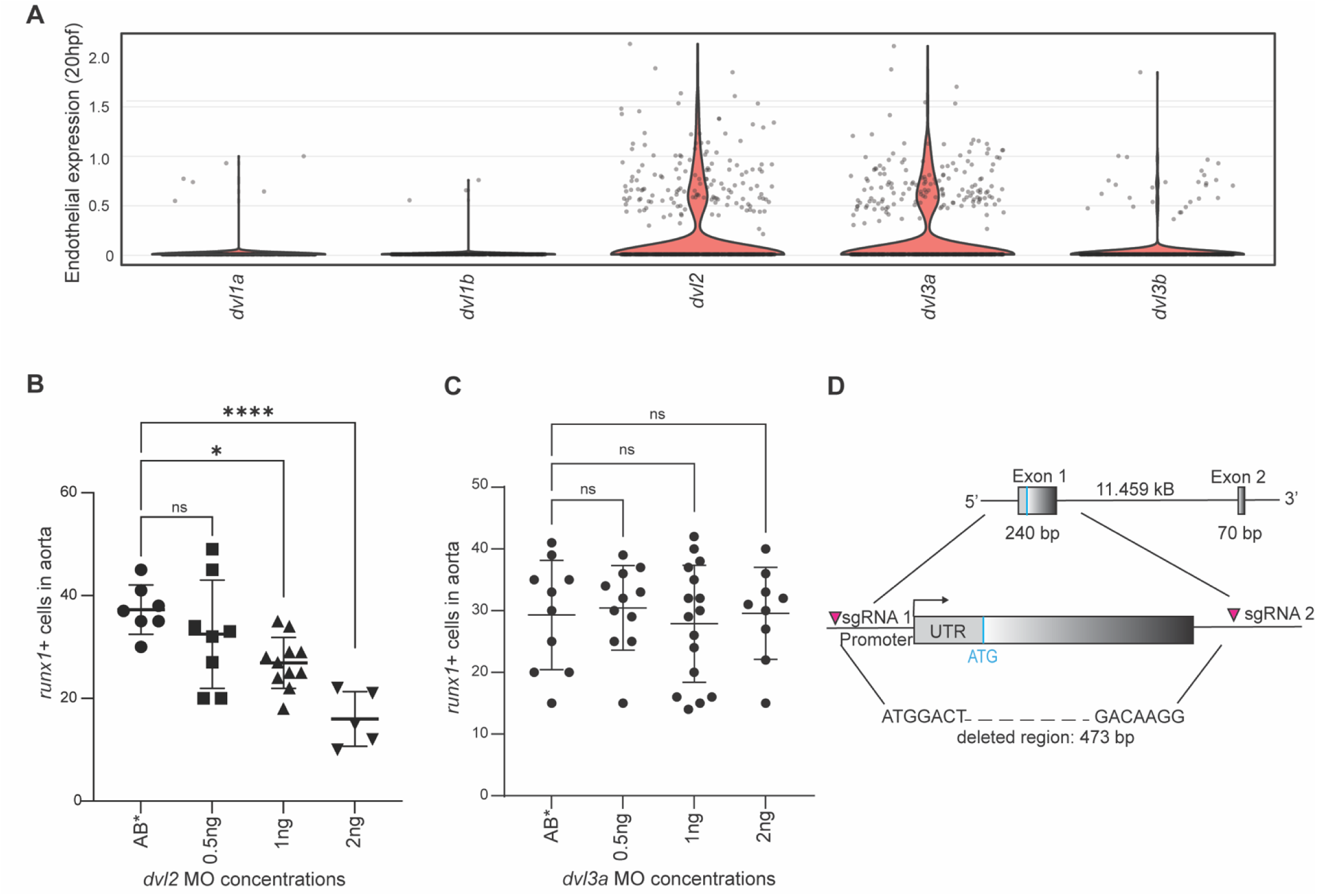
*dvl2* is expressed in endothelial cells and affects HSC development. **A**. Expression of all five *dvls* in endothelial cells at 20 hpf from DanioCell scRNA-Seq database (v1.0.3). **B.** Wild-type (AB*) embryos were injected with increasing concentrations (0.5ng/nL-2ng/nL) of *dvl2* splice blocking MO, fixed at 26 hpf and analyzed by WISH for *runx1*. Quantification of *runx1+* cells in the dorsal aorta with each dot representing a biological replicate. One-Way ANOVA with post hoc Tukey comparisons; * p<0.05, **** p<0.0001. **C**. Wild-type (AB*) embryos were injected with increasing concentrations (0.5ng/nL-2ng/nL) of *dvl3a* splice blocking MO, fixed at 26 hpf and analyzed by WISH for *runx1*. Quantification of *runx1+* cells in the dorsal aorta with each dot representing a biological replicate. One-Way ANOVA with post hoc Tukey comparisons. **D.** *dvl2* mutants were generated by injecting two guide RNAs targeting upstream of the promoter and ATG start site on exon 1.

**Supplementary Figure 2.**
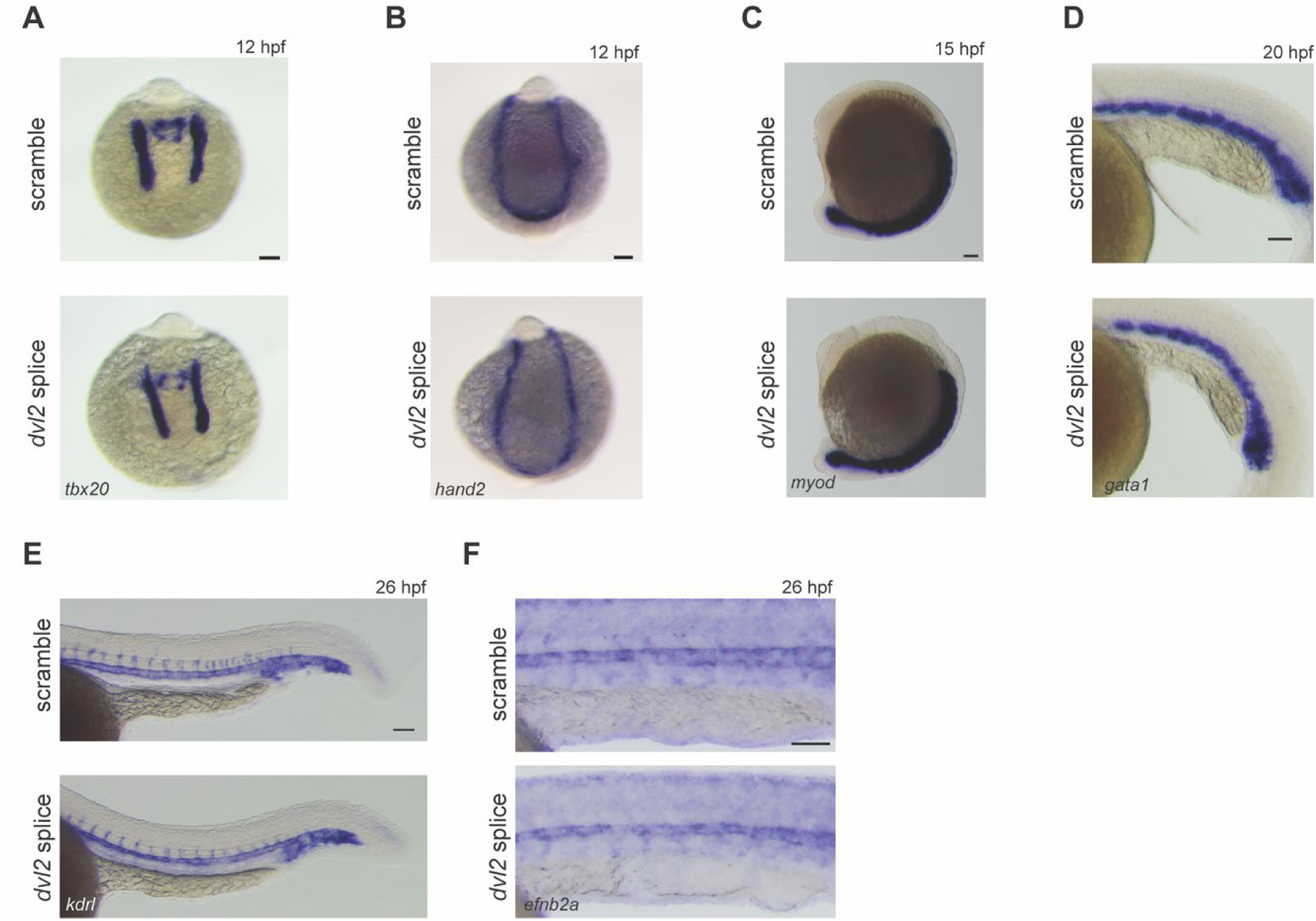
Dvl2 loss of function does not affect other developmental processes. Scramble MO or *dvl2* splice MO-injected embryos were analyzed by WISH for tissue-specific marker genes. Representative images are shown for **A.** *tbx20* and **B.** *hand2* (mesoderm), **C.** *myod* (somites), **D.** *gata1* (primitive blood), **E.** *kdrl* (endothelial cells), and **F.** *efnb2a* (arterial cells). Scale bar = 100 µm.

**Supplementary Figure 3.**
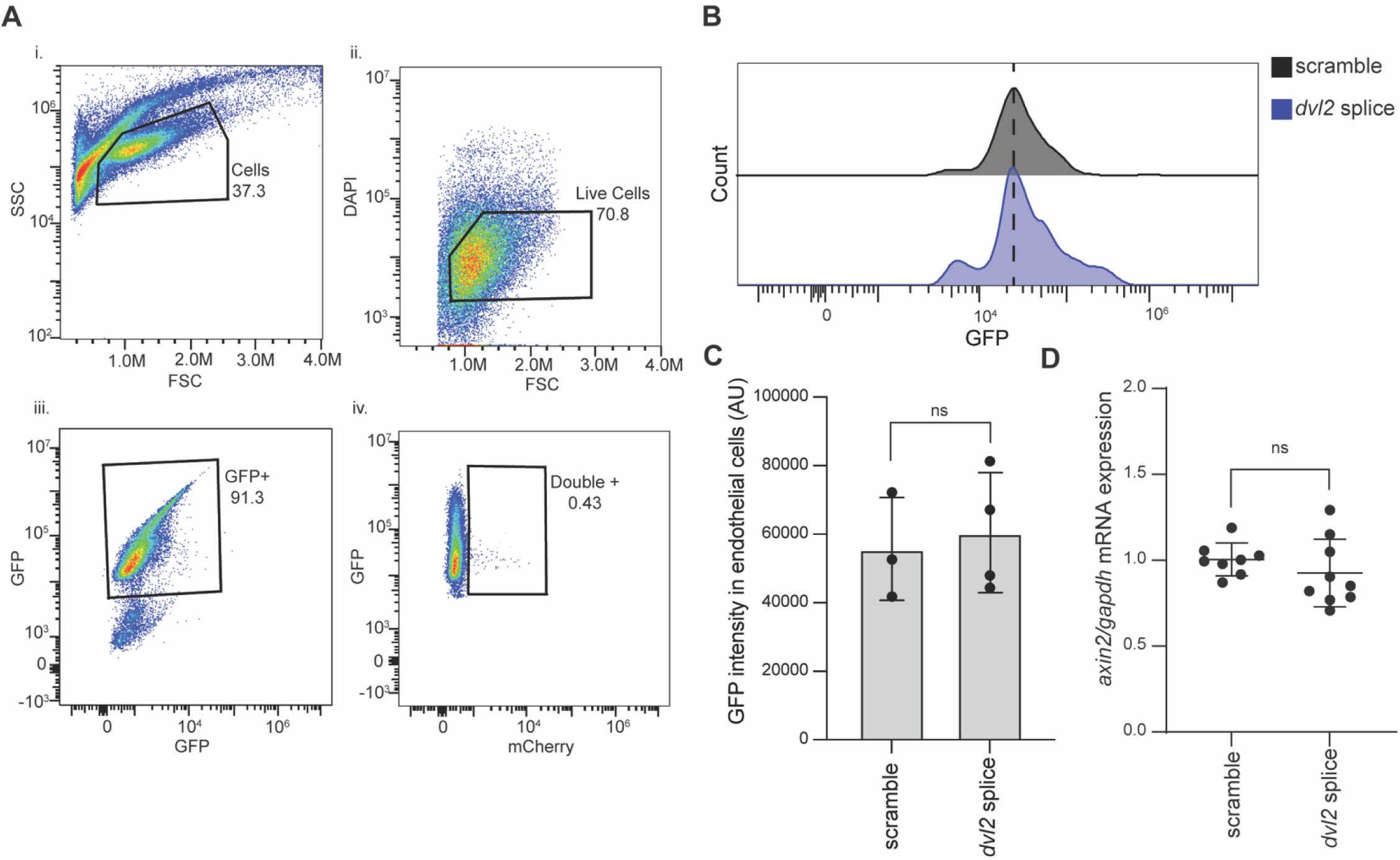
Loss of dvl2 does not affect canonical Wnt signaling. **A.** Flow cytometry gating strategy for *7XTCF:GFP;kdrl:NLS-mcherry* fish: i. Gating for cells, ii. Gating out dead cells using DAPI, iii. Gating for GFP+ cells, iv. Gating for double positive endothelial cells. B. Representative flow cytometry histogram of *7XTCF:GFP;kdrl:NLS-mcherry* fish injected with scramble MO or *dvl2* splice MO and the double positive cells were analyzed for GFP fluorescence at 16.5 hpf. Similar trends were observed in two additional trials. C. Flow cytometry quantification of GFP intensity in double positive cells (n= fish per biological replicate, 3 trials). Two-tailed Student’s. D. AB* embryos were injected with scramble MO or *dvl2* splice MO and expression of *axin2* was analyzed using qPCR. Two-tailed Student’s.

**Table 1.**
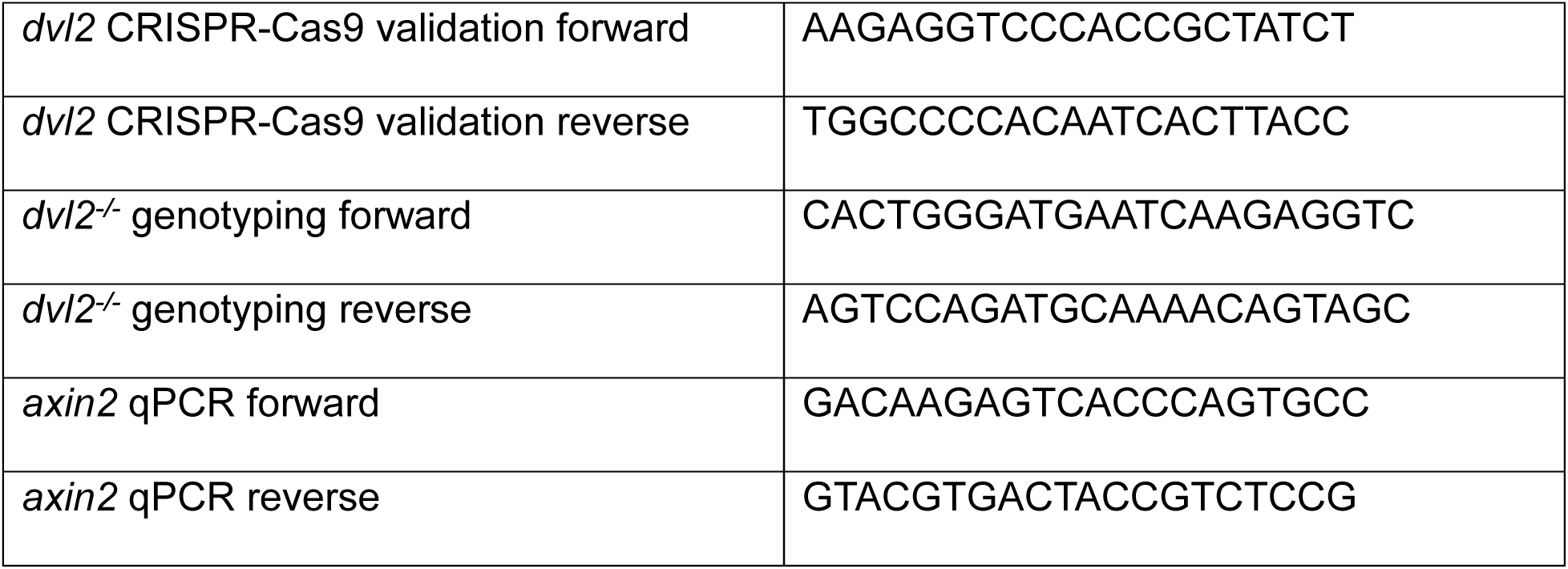
PCR and qPCR primers.

